# The itch-scratch reflex generates protective immunity

**DOI:** 10.1101/808477

**Authors:** Ali Radjavi

## Abstract

Itch: its complex neurobiology, its exquisite evolutionary conservation, and even the undeniably euphoric sensation of the scratch it evokes, are all suggestive of a productive physiological function. Nevertheless, we still struggle to answer (or altogether overlook) the basic question of why we itch in the first place. Here, we propose a simple hypothesis: the purpose of itch sensation is to evoke scratching behavior, which in turn boosts protective immunity against the broad range of pathogenic challenges that enter at the skin. We propose that the key function of itch induced scratching is to physically disrupt the skin, serving as a “mechanical adjuvant” that amplifies and directs immune responses to the precise site of potential pathogen entry. As proof of principle, we show that the potent adjuvanticity of itch inducing Compound 48/80 is dependent on this agent’s ability to elicit scratching behavior.

## Introduction

There’s a vague folk wisdom around the notion that you should not scratch an itch. Everything about that prescription is illustrated in a recent Readers Digest article entitled “Why you shouldn’t scratch a bug bite”^58^ in which “all experts agree” that scratching a bug bite is detrimental. Scratching can spread an infection, or leave the scratch site to intensely wounded that it could lead to a new infection. However, that article and so many like it, fail to acknowledge the rarity of these types of adverse outcomes from scratching a bug bite.

Our brains actually reward us for scratching. That sense of euphoria originates from dopaminergic neurons in the ventral tegmental area, the same reward center of the brain that’ give rise to the sensation of orgasm and is implicated in drug addiction^59^. So if scratching an itch is detrimental then why do our brains encourage us to do it? It *must* have a survival benefit, and whatever that is, it must greatly outweigh its possible adverse effects. Moreover that survival benefit must be broadly important to at least all warm blooded animals who have conserved the itch/scratch reflex throughout evolution.

We believe answer can be found in the common enemy of warm blooded animals; mosquitos, flies, lice, fleas and other biting pests (or more accurately to the pathogens and parasites that each of these pest carry). In humans alone these infections include malaria, dengue fever, yellow fever, west nile, zika, Japanese encephalitis, lymphatic filariasis. Lyme disease, tick-borne meningoencephalitis, Crimean–Congo hemorrhagic fever, tick-borne relapsing fever, Q fever, the tick-borne spotted fevers, babesiosis, ehrlichiosis, tularemia.Sleeping sickness, leishmaniasis, bartonellosis, relapsing fever, typhus, plague, and so on.

Many of these infections can kill you, *but* if you’re immune system can catch the infections early, you stand a much better chance of survival. And therein lies the dilemma. How can the immune system dedect an infection early, when the pathogens numbers are so few and the wound they came in through is so minuscule? The scratch reflex is the ingenious answer to this dilemma: let’s make the wound bigger and badder.

With a ubiquitous distribution of pathogen carrying biting arthropods that leave a non-feeling, often unperceivable bite of just a few microns on the nearly 2m^2^ surface of human skin, any mechanism that focuses and amplifies the immune response to pathogen deposited at the insult site would be tremendously advantageous. A large body of evidence has established the connection between mechanical disruption of skin tissue and inflammation. For instance, sterile scratching the mouse ear with a needle tip evokes neutrophil influx along the swath of the wound within 15 minutes of injury^1,2^, disruption of the stratum corneum by repeated tape strips stimulates antigen uptake by Langerhans cells^3^, even the remarkable efficacy of Jenner’s original smallpox vaccine came not from the immunogenicity of the strain as previously thought, but from unique method of inoculation–scarification of the skin^4^. While understandably dose dependent, we consider these laboratory models of superficial skin injury to partly recapitulate the natural scratch response elicited by itch. The same adjuvant like affect achieved in these methods could also be at play in the physiological itch/scratch response to insect bites–in a way that confers significant immunoprotection to skin intruding pathogens.

One previously proposed role for scratching behavior is to physically remove a biting pest from the skin^5^. The itch elicited by the landing of a mosquito or the crawling of a tick is an example of how this reflex may prevent insect bites in the first place. This type of touch-evoked itch likely operates through the excitement of low threshold mechanoreceptors of the skin, and is distinct from the itch elicited in response to the bite itself^6^. By contrast, a much more robust itch sensation develops after the biting insect has left the skin area, 10-15 minutes post bite in the case of mosquitos^7–9^, and on the order of hours and days in response to ticks^5^. We hypothesize that scratching as a result of this type of hypersensitivity-associated itch has less to do with removing the biting insect–which is probably long gone, but is meant to evoke self-inflicted skin disruption (i.e. scratching), which in turn localizes and amplifies the immune response to any potential pathogen delivered in the bite.

## The immunological significance of the skin stage of infection

The skin stage of infection is profoundly important to the efficacy of the anti-pathogen immune response. In the case of malaria, the significance of the early skin stage of infection has long been masked by models that infect mice intravenously with Plasmodium-infected red blood cells. Whereas the direct blood route achieves fulminant cerebral malaria in susceptible mice, the intradermal skin route of infection delivered through the mosquito vector appears to be immunologicaly protective^10,11^. The complex behaviors of plasmodium at the skin stage have only recently come to light^10–14^. Sporoozoites from the mosquito salivary gland are deposited in the skin, followed by migration to blood vessels in a process that can continue up to 2h post deposition^12,15,16^. Protective antigen specific CD8 T cells are primed in the lymph nodes that drain the skin bite^11^. Importantly many antigens are conserved between the developemental stages of Plasmodium^17^, making anti-sporozoite immunity protective against subsequent lifecycle stages^11^. Collectively, the skin stage of malaria infection may represent a short window of opportunity for immune activation prior to parasitization of the liver–an intrinsically immunosuppressive organ where Plasmodium is effectively cloaked from immunosurvailance^18^. While Plasmodium infection has been the most closely examined, the immunological significance of the skin stage of infection is likely a shared trait among all pathogens vectored by biting arthropods.

The newly appreciated importance of the skin stage of pathogenic infections, taken together with the immune activating outcome of mechanical disruption of the skin, lends attractiveness to the hypothesis that scratch may be operating as a “mechanical adjuvant”, boosting the local immune response and conferring meaningful immunoprotection.

## The immunomodulatory outcomes of skin barrier disruption

The elaborate immunological makeup of the skin is a testament to its importance as a barrier and sentinel of pathogenicity. Just beneath the uppermost stratum corneal layer of the undisturbed skin is a contiguous minefield of antigen-presenting Langerhans and dermal dendritic cells^19^. The same epidermal layer is punctuated with tissue resident T cells^20^, Mast cells^21^, and is continuously patrolled by circulating monocytes that extravasate through the vasculature and return to the circulation via skin draining lymphatics^22^. This elaborate but quiescent immune milieu quickly, and locally activates when the skin is mechanically disrupted.

Much of what we know about the immunological consequences of skin disruption comes from tape stripping models employed in allergy and contact hypersensitivity studies^1,2,23,24^. In response to just three tape strips, Kubo et al have visualized the elongation of Langerhan dendritic processes, which penetrate through the tight junctions of the keratinocyte layer above, where they can take up foreign antigen^24^. Amazingly, as Langerhans cells break up keratinocyte tight junctions, they concomitantly form new tight junctions around between their processes and the keratinocytes they contact, thereby never compromising the integrity of epidermis^24^. This study and several others have shown that within 24 hours of gentle tape stripping, Langerhans cells dramatically emigrate from the affected area and are recovered in the skin draining lymph node where they present captured antigen^3,24,25^. Dermal dendritic cells too upregulate markers of antigen presentation and migrate to regional lyphatics in response to skin barrier disruption^3^, and keratinocytes themselves have been shown to dramatically modulate their cytokine profiles in response to tape stripping^26^.

Not only does skin barrier disruption activate existing immune residents of the skin, but it leads to the immediate recruitment of circulating cells to the site of local disruption. Neutrophil influx occurs within minutes of tape striping or sterile skin scratch^1,2,27^as does the rapid infiltration of plasmacytoid dendritic cells^28^. While T cells are constitutively present in the skin, their numbers increase at 24 and peak at 48 hours post injury^28^, as did numbers of other immune players including innate lymphoid cells^29^, eusonophils^30^ and basophils^31^. Given the number of diverse cell types that respond to local skin disruption, one can foresee the diversity of accompanying changes in the local cytokine milieu^2,26–28,30,32^.

These immune responses to skin barrier disruption have profound influence on immune outcomes. The discussed upregulation of antigen presenting function and migration ultimately give rise to more robust T and B cell responses. Tape striping of the skin has even been used as an adjuvant to prime tumor specific cytotoxic T lymphocytes^33^. Not only does skin barrier disruption affect the magnitude of the adaptive immune response, but it also affects its nature and quality. Tape stripping experiments have clearly been shown to shift the TH1/TH2 balance of the adaptive immune response–though the direction of that shift is not absolute. Several studies associate scratching and tape-stripping with exacerbation of TH2 biased allergic immunity^30,34,35^, while other studies evidence a TH1 skewing effect of tape striping^28,36^. So while it’s clear that the TH1/TH2 balance is altered via skin disruption, and that this balance is critical for disease outcomes, both relationships appear to be complex, model, and timepoint dependent.

## The itch response is generated by the immune system

Most of the aforementioned studies have looked at the response to skin barrier disruption inflicted for the first time in a naïve animal. However in the context a mosquito or tick bite, an itch sensation actually implies a secondary bite exposure–i.e. type 1 hypersensitivity.

A study by Ohtsuka et al describes the development of a robust scratch response only after repeated mosquito bites over a sensitization interval ^37^. This study as well as others long before it^7,8,19^, suggest that there is nothing immediately pruritogenic in the mosquito salivary gland, but rather that the itch response is an acquired hypersensitivity reaction that is dependent on a period of sensitization. Importantly, we suggest that the onset of itch is a functional outcome of sensitization, empowering the skin to illicit itch sensation upon a subsequent bite challenge.

It is interesting to note that presensitization of mice with bites from uninfected mosquito’s conferred significant immunoprotection against subsequent bites from plasmodium infected mosquitos^14,38^. Similarly, presensitization with uninfected tick and sandfly bites appears to be protective against challenge with Borrelia and Leishmania infected ticks and sandflys respectively^5,39–42^. Remarkably, human patients who specifically reported itching in association with tick bites were less likely to develop lyme disease relative to non-itch reporting individuals^5^. We hypothesize that the immunoprotection reported in these models is scratch dependent and based on our proposed mechanism of mechanical adjuvanticity.

## Itch sensation: a neuro-immune interface

Ultimately the sensation of itch (and the reflex to scratch) is the thing of neurons. While the itch specific MrgprA3^+^ fibers that innervate the skin extend just beneath the stratum granulosum^43^, they do not sense itch directly. Instead, innervating itch fibers can be activated by a host of endogenous signaling molecules of the immune system^44^. Perhaps the best understood system is the histamine dependent itch pathway, in which antigen challenge at the skin results in degranulation of histamine loaded mast cells activated by antigen-IgE complex (with IgE generated from a prior sensitization event)^45^. As it relates to itch, histamine has been shown to activate itch specific, mechanically insensitive C fibers which in turn transmit signal to the spinal chord and brain to elicit scratching behavior^44^. Moreover, scratching is understood to activate pain receptors, which in turn inhibit itch receptor signaling and alleviate itch sensation^44,46^.

While histamine is the best-appreciated endogenous activator of itch, it is not necessary for itch induction as it is only one of many endogenous itch-inducing signals. In fact, a histamine receptor antagonist that blocks histamine induced itch failed to inhibit scratching behavior in mosquito bite sensitized mice, and mast cell deficient mice did not significantly diminish scratching behavior in response to mosquito bites^37^. TRPA1^47^, TLR7 agonsists^48^, TLR-3 agonists^49^, TSLP^50^ and IL-31^51^, have all been reported to induce itch activation. This apparent redundancy in the endogenous immune activators (and neuronal receptors) of itch, intimates not only the immunological importance of the itch/scratch reflex, but perhaps also to a hidden arms race between the immune system and skin-entering pathogens–aiming respectively, to induce, and suppress itch sensation.

## The itch inducing compound 48/80 has immune activating properties that are dependent on scratch

The OTII mouse was a practical and available choice for demonstrating proof of principle for our model. OTII mice have a nearly monoclonal T cell repertoire directed against irrelevant chicken egg ovalbumin (ova)^52^. When OTII mice are intradermaly challenged with ova, T cell activation can easily be measured–owing to the artificially high occurrence of ova-reactive T cells in the animal. In WT mice challenged with ova, antigen specific T cell activation is not readily measurable without the use of CD4 tetramers, owing to the naturally low clonal abundance of ova-reactive T cells^53^.

We injected OTII mice intradermaly, on the left cheek with non-endotoxin free ova (Sigma), alone or in combination with the itch-inducing compound 48/80 (Sigma). 24h later we harvested the draining brachial lymph nodes, and stained single cell suspensions for the CD4 T cell marker and the CD69 early T cell activation marker. Compound 48/80, potentiated higher CD69 expression on T cells than ova alone (Figure 1a). This adjuvant-like property was not limited to CD4 T cells in OTII mice, as wildtype C57BL/6 mice similarly challenged with ova+compound 48/80, showed higher titers of anti ova IgG1 and IgG2b than mice challenged with ova alone (Figure 1b).

**Figure 1.**
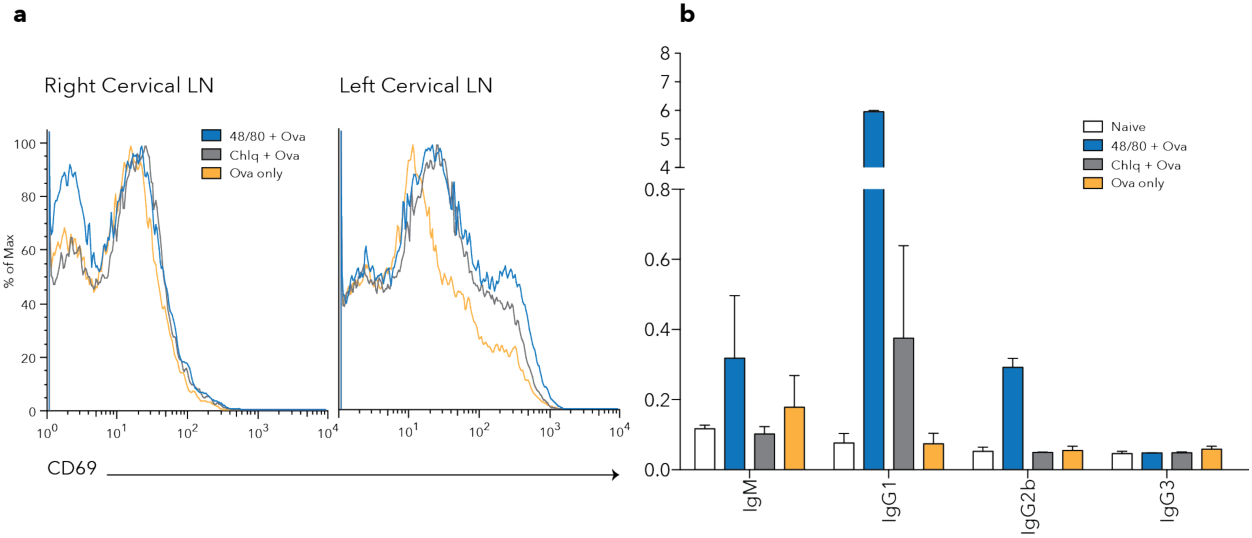
Itch inducing compounds enhance early T cell activation and antibody production. **a** Representative FACS analyses of early activation marker CD69 in cervical lymph nodes of OTII mice immunizec with ovalbumin at the left cheek, with or without chloroquin or compounc 48/80 prurirogens. 24h post immunization, **b** OD450 of ovalbumin specific antibody isotypes in WT mice 18 days after immunization with ovalbumin with or without chloroquin or compounc 48/80 pruritogens

Would the adjuventicity of compound 48/80 be lost if scratching was physically precluded? To test whether adjuvant like properties of compound 48/80 is derived from its ability to elicit scratching behavior, we needed to physically block skin scratching. To this end, we found commercially available jackets for mice to be inadequate. Instead, we developed our own custom jackets with a plastic guard that effectively protected against scratching at the left shoulder (Figure 2a). To control for any effect of discomfort or restraint owing to the plastic scratch guard, control “scratch permissive” jackets were made with the same plastic flap on the unimmunized right shoulder. Inhibiting scratching at the site of ova injection, effectively blocked the CD69 and anti-ova IgG1 activating property of compound 48/80 (Figure 2b), demonstrating that the adjuvanicity of compound 48/80 is derived from its ability to evoke scratching. An identical loss of effect was observed when ova+compound 48/80 was administered at the left cheek and scratching was inhibited using a cone shaped Elizabethan collar. While both modes of scratch inhibition yielded a loss of the adjuvant like effect of compound 48/80, only the jackets could be said to control for the confounding effects of psychological stress on the immune response^54–56^.

**Figure 2.**
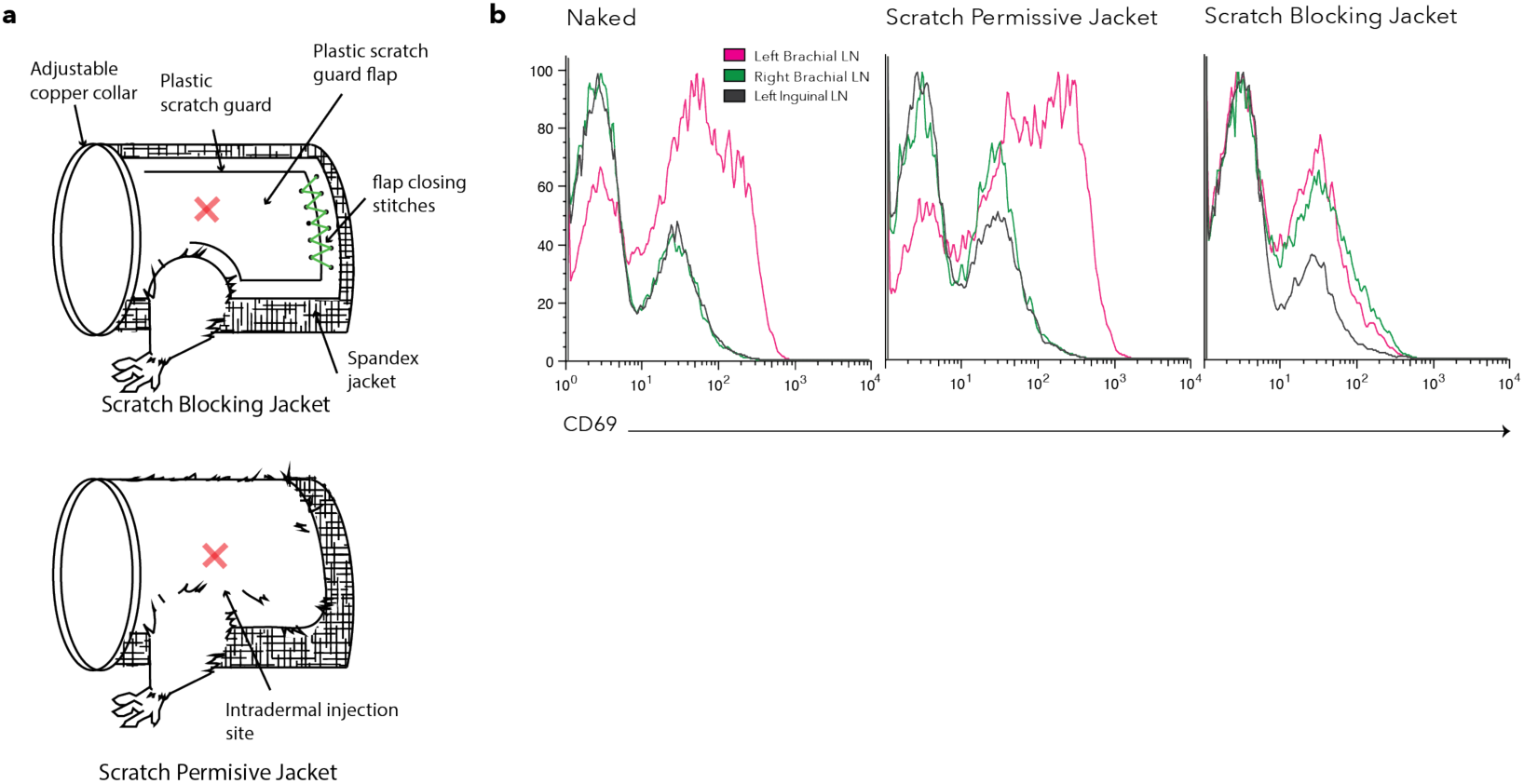
The adjuvant properties of itch inducing compound 48/80 are scratch dependent. **a.** Scratch blocking and scratch permissive mouse jackets, **b**, OTII mice were untouched (left), fitted with scratch permissivejackets (middle), or scratch blocking jackets (right), and immunized with ovalbumin+ compound 48/80 at the left shoulder. Representative-FAGS analyses of early activation marker CD69 in shoulder draining brachial lymph nodes, and control inguinal lymph node of each group 24h post immunization.

This simple model, while admittedly artificial, demonstrates proof of principle; Itch sensation at a site of antigen delivery can give rise to more robust early T cell activation, and that this effect is entirely dependent on physical scratching of the insult site. Indeed, the adjuvant like properties of compound 48/80 have been previously reported^57^, however we believe we are the first to show that its adjuvanticity is likely not an intrinsic property of the compound, but is dependent on its evocation of scratching behavior.

## Conclusions

It is clear that the immunological consequence of skin barrier disruption does not operate on a single cell type or cytokine, but modulates virtually every part of the skins immune repertoire. Accordingly, our central hypothesis that itch evoked scratching behavior is immunoprotective, likely does not operate via a single mechanism, but rather through the collective contribution of each aforementioned consequence of skin barrier disruption, including antigen uptake, APC migration, neutrophil/basophil/eusonophil/pDCs/ILC/influx, T cell activation and skewing, B cell isotype switching and cytokine modulation, all of which can impact the efficacy and outcome of an immune response to skin-penetrating pathogens.

Importantly, the extensive immunomodulatory effects of skin barrier disruption, give rise to numerous points of possible deregulation, making the association between scratching and allergy not a surprising one^23^. Indeed, one hope is that in appreciating the normal physiological function of the itch/scratch reflex we might provide an invaluable context for our understanding of itch related skin and neurological pathologies.

While mosquito bites and plasmodium infection is a heavily referenced example of itch induction and anti pathogen immunity, a protective role for the itch scratch reflex based on a model of mechanical ajduvanticity represents a line of defense that is applicable to the astounding list of pathogens vectored by biting arthropods. When the morbidity of these infections is taken into consideration, the evolutionary pressure to generate, conserve, and reward what we propose here to be a protective reflex, is immediately clear. If our hypothesis is correct, then considering the incalculable number of mosquito, tick, mite and sandfly bites a year, public knowledge of a robust protective function for the itch/scratch reflex may confer meaningful and cost-free protection from the host of blood-borne pathogens vectored by parasitic arthropods.

## References

1. Ng, L. G. et al. Visualizing the neutrophil response to sterile tissue injury in mouse dermis reveals a three-phase cascade of events. J. Invest. Dermatol. 131, 2058– 2068 (2011).

2. Oyoshi, M. K. et al. Leukotriene B4-driven neutrophil recruitment to the skin is essential for allergic skin inflammation. Immunity 37, 747–758 (2012).

3. Kissenpfennig, A. et al. Dynamics and function of Langerhans cells in vivo: dermal dendritic cells colonize lymph node areas distinct from slower migrating Langerhans cells. Immunity 22, 643–654 (2005).

4. Liu, L. et al. Epidermal injury and infection during poxvirus immunization is crucial for the generation of highly protective T cell-mediated immunity. Nat. Med. 16, 224–227 (2010).

5. Burke, G. et al. Hypersensitivity to Ticks and Lyme Disease Risk. Emerg. Infect. Dis. 11, 36–41 (2005).

6. Akiyama, T. & Carstens, E. Neural processing of itch. Neuroscience 250, 697–714 (2013).

7. McKiel, J. A. Sensitization to Mosquito Bites. Can. J. Zool. 37, 341–351 (1959).

8. Wilson, A. B. & Clements, A. N. THE NATURE OF THE SKIN REACTION TO MOSQUITO BITES IN LABORATORY ANIMALS. Int. Arch. Allergy Appl. Immunol. 26, 294–314 (1965).

9. Reunala, T., Brummer-Korvenkontio, H. & Palosuo, T. Are we really allergic to mosquito bites? Ann. Med. 26, 301–306 (1994).

10. Spence, P. J. et al. Vector transmission regulates immune control of Plasmodium virulence. Nature 498, 228–231 (2013).

11. Chakravarty, S. et al. CD8+ T lymphocytes protective against malaria liver stages are primed in skin-draining lymph nodes. Nat. Med. 13, 1035–1041 (2007).

12. Sinnis, P. & Zavala, F. The skin: where malaria infection and the host immune response begin. Semin. Immunopathol. 34, 787–792 (2012).

13. Guilbride, D. L., Guilbride, P. D. L. & Gawlinski, P. Malaria’s deadly secret: a skin stage. Trends Parasitol. 28, 142–150 (2012).

14. Kebaier, C., Voza, T. & Vanderberg, J. Kinetics of mosquito-injected Plasmodium sporozoites in mice: fewer sporozoites are injected into sporozoite-immunized mice. PLoS Pathog. 5, e1000399 (2009).

15. Sidjanski, S. & Vanderberg, J. P. Delayed migration of Plasmodium sporozoites from the mosquito bite site to the blood. Am. J. Trop. Med. Hyg. 57, 426–429 (1997).

16. Matsuoka, H., Yoshida, S., Hirai, M. & Ishii, A. A rodent malaria, Plasmodium berghei, is experimentally transmitted to mice by merely probing of infective mosquito, Anopheles stephensi. Parasitol. Int. 51, 17–23 (2002).

17. Hollingdale, M. R. & Krzych, U. Immune responses to liver-stage parasites: implications for vaccine development. Chem. Immunol. 80, 97–124 (2002).

18. Knolle, P. A. & Gerken, G. Local control of the immune response in the liver. Immunol. Rev. 174, 21–34 (2000).

19. Bos, J. D. & Kapsenberg, M. L. The skin immune system Its cellular constituents and their interactions. Immunol. Today 7, 235–240 (1986).

20. Carbone, F. R. & Gebhardt, T. Immunology. A neighborhood watch upholds local immune protection. Science 346, 40–41 (2014).

21. Galli, S. J., Grimbaldeston, M. & Tsai, M. Immunomodulatory mast cells: negative, as well as positive, regulators of immunity. Nat. Rev. Immunol. 8, 478– 486 (2008).

22. Mueller, S. N., Gebhardt, T., Carbone, F. R. & Heath, W. R. Memory T cell subsets, migration patterns, and tissue residence. Annu. Rev. Immunol. 31, 137– 161 (2013).

23. De Benedetto, A., Kubo, A. & Beck, L. A. Skin barrier disruption: a requirement for allergen sensitization? J. Invest. Dermatol. 132, 949–963 (2012).

24. Kubo, A., Nagao, K., Yokouchi, M., Sasaki, H. & Amagai, M. External antigen uptake by Langerhans cells with reorganization of epidermal tight junction barriers. J. Exp. Med. 206, 2937–2946 (2009).

25. Holzmann, S. et al. A model system using tape stripping for characterization of Langerhans cell-precursors in vivo. J. Invest. Dermatol. 122, 1165–1174 (2004).

26. Nickoloff, B. J. & Naidu, Y. Perturbation of epidermal barrier function correlates with initiation of cytokine cascade in human skin. J. Am. Acad. Dermatol. 30, 535–546 (1994).

27. Keijsers, R. R. M. C. et al. In vivo induction of cutaneous inflammation results in the accumulation of extracellular trap-forming neutrophils expressing RORγt and IL-17. J. Invest. Dermatol. 134, 1276–1284 (2014).

28. Gregorio, J. et al. Plasmacytoid dendritic cells sense skin injury and promote wound healing through type I interferons. J. Exp. Med. 207, 2921–2930 (2010).

29. Kim, B. S. Innate lymphoid cells in the skin. J. Invest. Dermatol. 135, 673–678 (2015).

30. Onoue, A., Kabashima, K., Kobayashi, M., Mori, T. & Tokura, Y. Induction of eosinophil- and Th2-attracting epidermal chemokines and cutaneous late-phase reaction in tape-stripped skin. Exp. Dermatol. 18, 1036–1043 (2009).

31. Cheng, L. E. et al. IgE-activated basophils regulate eosinophil tissue entry by modulating endothelial function. J. Exp. Med. 212, 513–524 (2015).

32. Wood, L. C. et al. Barrier disruption increases gene expression of cytokines and the 55 kD TNF receptor in murine skin. Exp. Dermatol. 6, 98–104 (1997).

33. Seo, N. et al. Percutaneous peptide immunization via corneum barrier-disrupted murine skin for experimental tumor immunoprophylaxis. Proc. Natl. Acad. Sci. U. S. A. 97, 371–376 (2000).

34. Oyoshi, M. K., Larson, R. P., Ziegler, S. F. & Geha, R. S. Mechanical injury polarizes skin dendritic cells to elicit a Th2 response by inducing cutaneous TSLP expression. J. Allergy Clin. Immunol. 126, 976–984.e5 (2010).

35. Strid, J., Callard, R. & Strobel, S. Epicutaneous immunization converts subsequent and established antigen-specific T helper type 1 (Th1) to Th2-type responses. Immunology 119, 27–35 (2006).

36. Matsushima, H., Hayashi, S. & Shimada, S. Skin scratching switches immune responses from Th2 to Th1 type in epicutaneously immunized mice. J. Dermatol. Sci. 32, 223–230 (2003).

37. Ohtsuka, E. et al. Roles of mast cells and histamine in mosquito bite-induced allergic itch-associated responses in mice. Jpn. J. Pharmacol. 86, 97–105 (2001).

38. Donovan, M. J. et al. Uninfected mosquito bites confer protection against infection with malaria parasites. Infect. Immun. 75, 2523–2530 (2007).

39. Wikel, S. K., Ramachandra, R. N., Bergman, D. K., Burkot, T. R. & Piesman, J. Infestation with pathogen-free nymphs of the tick Ixodes scapularis induces host resistance to transmission of Borrelia burgdorferi by ticks. Infect. Immun. 65, 335–338 (1997).

40. Nazario, S. et al. Prevention of Borrelia burgdorferi transmission in guinea pigs by tick immunity. Am. J. Trop. Med. Hyg. 58, 780–785 (1998).

41. Gomes, R. B. et al. Seroconversion against Lutzomyia longipalpis saliva concurrent with the development of anti-Leishmania chagasi delayed-type hypersensitivity. J. Infect. Dis. 186, 1530–1534 (2002).

42. Valenzuela, J. G. et al. Toward a defined anti-Leishmania vaccine targeting vector antigens: characterization of a protective salivary protein. J. Exp. Med. 194, 331–342 (2001).

43. Han, L. et al. A subpopulation of nociceptors specifically linked to itch. Nat. Neurosci. 16, 174–182 (2013).

44. Bautista, D. M., Wilson, S. R. & Hoon, M. A. Why we scratch an itch: the molecules, cells and circuits of itch. Nat. Neurosci. 17, 175–182 (2014).

45. Galli, S. J. & Tsai, M. IgE and mast cells in allergic disease. Nat. Med. 18, 693– 704 (2012).

46. Davidson, S., Zhang, X., Khasabov, S. G., Simone, D. A. & Giesler, G. J. Relief of itch by scratching: state-dependent inhibition of primate spinothalamic tract neurons. Nat. Neurosci. 12, 544–546 (2009).

47. Wilson, S. R. et al. TRPA1 is required for histamine-independent, Mas-related G protein-coupled receptor-mediated itch. Nat. Neurosci. 14, 595–602 (2011).

48. Liu, T., Xu, Z.-Z., Park, C.-K., Berta, T. & Ji, R.-R. Toll-like receptor 7 mediates pruritus. Nat. Neurosci. 13, 1460–1462 (2010).

49. Liu, T. et al. TLR3 deficiency impairs spinal cord synaptic transmission, central sensitization, and pruritus in mice. J. Clin. Invest. 122, 2195–2207 (2012).

50. Wilson, S. R. et al. The epithelial cell-derived atopic dermatitis cytokine TSLP activates neurons to induce itch. Cell 155, 285–295 (2013).

51. Grimstad, O. et al. Anti-interleukin-31-antibodies ameliorate scratching behaviour in NC/Nga mice: a model of atopic dermatitis. Exp. Dermatol. 18, 35–43 (2009).

52. Barnden, M. J., Allison, J., Heath, W. R. & Carbone, F. R. Defective TCR expression in transgenic mice constructed using cDNA-based alpha- and beta-chain genes under the control of heterologous regulatory elements. Immunol. Cell Biol. 76, 34–40 (1998).

53. Hataye, J., Moon, J. J., Khoruts, A., Reilly, C. & Jenkins, M. K. Naïve and Memory CD4+ T Cell Survival Controlled by Clonal Abundance. Science 312, 114–116 (2006).

54. Dhabhar, F. S., Malarkey, W. B., Neri, E. & McEwen, B. S. Stress-induced redistribution of immune cells--from barracks to boulevards to battlefields: a tale of three hormones--Curt Richter Award winner. Psychoneuroendocrinology 37, 1345–1368 (2012).

55. Dhabhar, F. S., Miller, A. H., McEwen, B. S. & Spencer, R. L. Effects of stress on immune cell distribution. Dynamics and hormonal mechanisms. J. Immunol. Baltim. Md 1950 154, 5511–5527 (1995).

56. Saint-Mezard, P. et al. Psychological stress exerts an adjuvant effect on skin dendritic cell functions in vivo. J. Immunol. Baltim. Md 1950 171, 4073–4080 (2003).

57. McGowen, A. L., Hale, L. P., Shelburne, C. P., Abraham, S. N. & Staats, H. F. The mast cell activator compound 48/80 is safe and effective when used as an adjuvant for intradermal immunization with Bacillus anthracis protective antigen. Vaccine 27, 3544–3552 (2009).

58. Cahn L 2017 ‘Why you shouldn’t scratch a bug bite’, Readers Digest, 19 July, accessed October 2017

59. Su, X.Y., Chen, M., Yuan, Y., Li, Y., Guo, S.S., Luo, H.Q., Huang, C., Sun, W., Li, Y., Z hu, M.X., et al Central Processing of Itch in the Midbrain Reward Center. Neuron (2019).

